# Diurnal variation of brain activity in the human suprachiasmatic nucleus

**DOI:** 10.1101/2023.07.10.548316

**Authors:** Satoshi Oka, Akitoshi Ogawa, Takahiro Osada, Masaki Tanaka, Koji Nakajima, Koji Kamagata, Shigeki Aoki, Yasushi Oshima, Sakae Tanaka, Eiji Kirino, Takahiro J. Nakamura, Seiki Konishi

## Abstract

The suprachiasmatic nucleus (SCN) is the central clock for circadian rhythms. Animal studies have revealed daily rhythms in the neuronal activity in the SCN. However, the circadian activity of the human SCN has remained elusive. In this study, to reveal the diurnal variation of the SCN activity in humans, the SCN was localized, and its activity was investigated using perfusion imaging. We scanned each participant four times a day, every six hours, and higher activity was observed at noon while lower activity was recorded in the early morning. The SCN activity was then measured every thirty minutes for six hours from midnight to dawn and showed a decreasing trend and was comparable with the rodent SCN activity after switching off the lights. These results suggest that the diurnal variation of the human SCN follows the zeitgeber cycles of mammals and is modulated by physical lights rather than the local time.

## Introduction

Circadian rhythms control physiology and behavior with the 24-hour cycle of day and night^1^ and circadian disruption is related to diseases in various human systems, including autonomic and endocrine systems^2, 3^. The suprachiasmatic nucleus (SCN) is the central clock for circadian rhythms in mammals^4, 6–16, 51^. The SCN is a pair of nuclei in the anterior part of the hypothalamus, located above the optic chiasm and lateral to the third ventricle. Intrinsically photosensitive retinal ganglion cells that express the photopigment, melanopsin, project to the SCN neurons via the retinohypothalamic tract ^17–20^. Modulated by light exposure, the SCN activity is the master clock for the body clock cycles in animals. The SCN sends the clock information to other parts of the central nervous system and influences the autonomic and endocrine systems. Moreover, the SCN affects motor activity^21^, emotion^22, 23^, and cognition such as memory^18, 23–25^.

Several previous studies have shown some basic features of the SCN in humans. Postmortem brain studies showed diurnal cycles of neuropeptides in the human SCN^26–29^. In functional magnetic resonance imaging (fMRI), it has been shown that visual responses of the SCN depend on the light wavelength^29^. In contrast to animal studies, it is hard to conduct experiments in humans due to the difficulties in strictly controlling the duration of light and darkness. It is also important to note that modern lifestyle influences the internal clock, such as excessive use of bright light until late at night^20^. Moreover, because of the small size of the SCN, measuring the SCN activity in humans remained difficult. However, how the SCN activity changes over one cycle of the circadian rhythm and how the circadian SCN activity differs from those in animals need to be investigated. This study investigated the diurnal variation of the human SCN activity using perfusion imaging that measures cerebral blood flow (CBF) in the brain.

Before the scan, we localized the SCN by using the resting state images collected in our previous study^30^. Brain activity in the SCN over 24 hours was then measured. The participants were scanned four times a day, every six hours, with a within-participant design. Brain activity was further scanned every thirty minutes for six hours from midnight to dawn to reveal more detailed characteristics of the diurnal variation compared to previous animal data.

## Results

Two experiments were conducted in this study. In the first experiment, the whole cycle of the diurnal activity of the SCN was investigated by two perfusion images measuring the CBF, which were acquired four times (18:00, 24:00, 6:00, 12:00 on local time) within 24 hours (Experiment 1) (Fig. 1a, Supplementary Fig. 1). The lights in the MRI room were on in all the scans. On the other hand, the second experiment investigated the temporal trend of human SCN activity at night (Experiment 2). The perfusion images were scanned every 30 min from 24:00 to 6:00 on local time (Fig. 1b), with electroencephalography (EEG) recorded using an MRI-compatible system. The lights of MRI room were switched off at 0:15 and on at 5:45. To analyze the CBF in the SCN, the SCN was localized by using the blood-oxygenation-level-dependent (BOLD) images during the resting state collected in our previous study^30^. An areal boundary mapping technique parcellated the anterior part of the hypothalamus into the subregions, including the SCN.

**Fig. 1.**
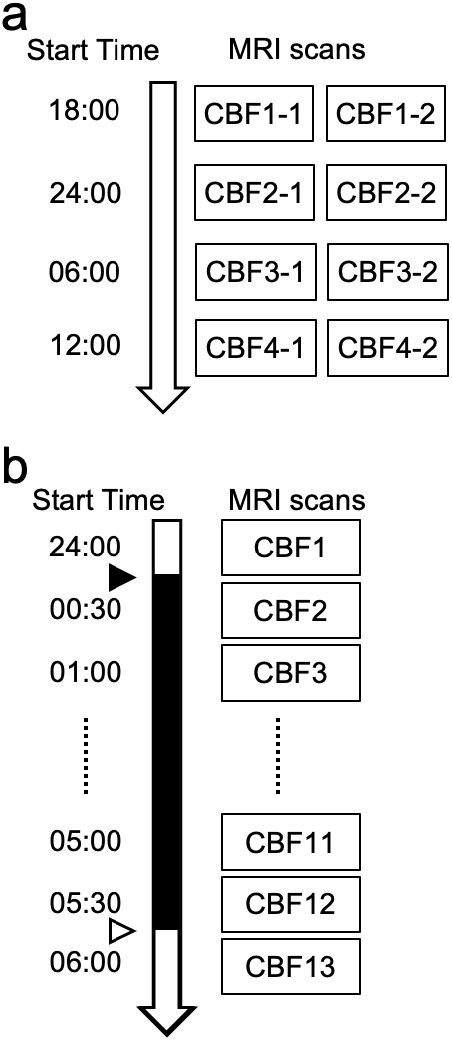
Scan schedules. **a** Scan schedule of Experiment 1. Scans were conducted in evening (18:00), midnight (24:00), morning (6:00), and daytime (12:00) sessions in that order. Two CBF images were acquired and averaged at each session to increase the signal-to-noise ratio. Between the scan sessions, the participants rested at the accommodation provided. **b** Scan schedule of Experiment 2. The room lights were on during the first scan (called pre) and the last scan (called post), whereas the room lights were off during the scans from 0:15 to 5:45. The black and white arrowheads indicate the lights in the MRI room being switched off and on, respectively. The black bar indicates the period in which the light of MRI room was off.

### SCN localization

The areal boundary mapping technique generated, from BOLD images^30^, the probabilistic boundary map in 1.25-mm resolution, where the probability value represented how likely the voxel is a boundary of a functional area, a “parcel” that corresponds to a hypothalamic nucleus (Fig. 2a/b). The low probability value in a voxel of the map, therefore, indicates that the voxel is likely the center of a parcel. In the probability map, the SCN was identified bilaterally above the optic chiasm and lateral to the third ventricle (Supplementary Fig. 2a/b). We also localized the surrounding nuclei and listed the coordinates in Table 1. The coordinates of the voxel of the SCN ROI in 2- mm resolution for perfusion imaging were x = -2, y = 2, z = -16 for the left ROI and x = 2, y = 2, z = -16 for the right ROI (Fig. 2a/b).

**Fig. 2.**
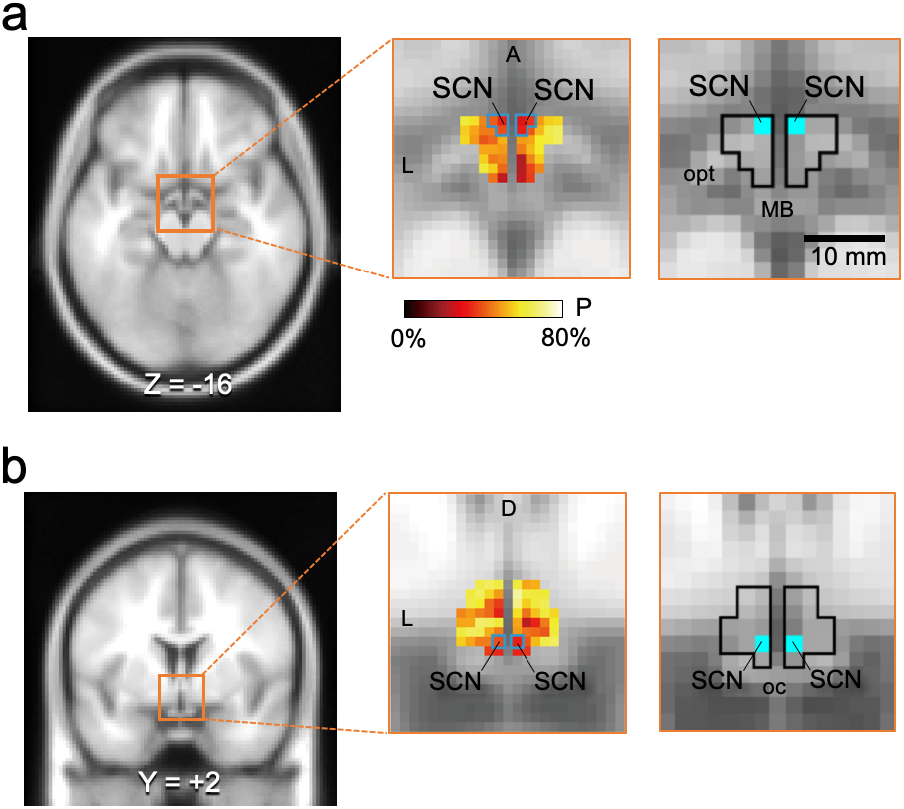
ROI definition of the SCN. **a** SCN ROI on the axial section. The probabilistic boundary map of the hypothalamus is shown on the axial section (center panel). High probability (yellow) indicates that the boundaries are likely to exist, while lower probability than the surroundings (red) indicates that the centers of nuclei are likely to exist. The right panel shows the SCN ROI on the axial section. **b** ROIs of the SCN on the coronal section. The center panel shows the probabilistic boundary map, while the right panel shows the ROIs of the SCN. Black lines indicate the border of the hypothalamus. L = left, R = right, D = dorsal, V = ventral, A = anterior, oc = optic chiasma, opt = optic tract, MB = mammillary body, 3V = third ventricle.

**Table 1.** Centroid coordinates of the SCN and neighboring hypothalamic nuclei and their distances from the SCN.

| Nucleus | *x* | *y* | *z* | Distance from SCN (mm) |
| --- | --- | --- | --- | --- |
| L SCN | -1.8 | +1.5 | -15.8 | - |
| R SCN | +1.8 | +1.5 | -15.8 | - |
| L MPO | -1.5 | +2.3 | -12.8 | 3.3 |
| R MPO | +1.5 | +2.3 | -12.8 | 3.3 |
| L SO | -5.5 | +1.8 | -12.8 | 4.8 |
| R SO | +4.3 | +1.0 | -12.5 | 4.2 |
| L PVH | -1.8 | -0.5 | -10.5 | 5.7 |
| R PVH | +1.5 | +0.3 | -10.8 | 5.2 |

L = left, R = right, SO = supraoptic nucleus, PVH = paraventricular nucleus of the hypothalamus.

### Diurnal variation of SCN activity

Experiment 1 investigated whether the SCN activity had increased or decreased signal changes within 24 hours. We performed a one-way repeated-measures analysis of variance (ANOVA) over the time (6:00, 12:00, 18:00, and 24:00 on local time). The activity in the SCN was highest at 12:00 in the daytime and lowest at 6:00 in the morning (Fig. 3a). The SCN activity was significantly modulated among the four scans (*F*(3,78) = 3.38, *P* = 0.022). The activity at 12:00 was significantly higher than that at 6:00 (Tukey-Kramer test, *P* < 0.05). A similar trend was seen when the left and right SCN were analyzed separately (Fig. 3a) and when the first and second scans were analyzed separately (Fig. 3b).

**Fig. 3.**
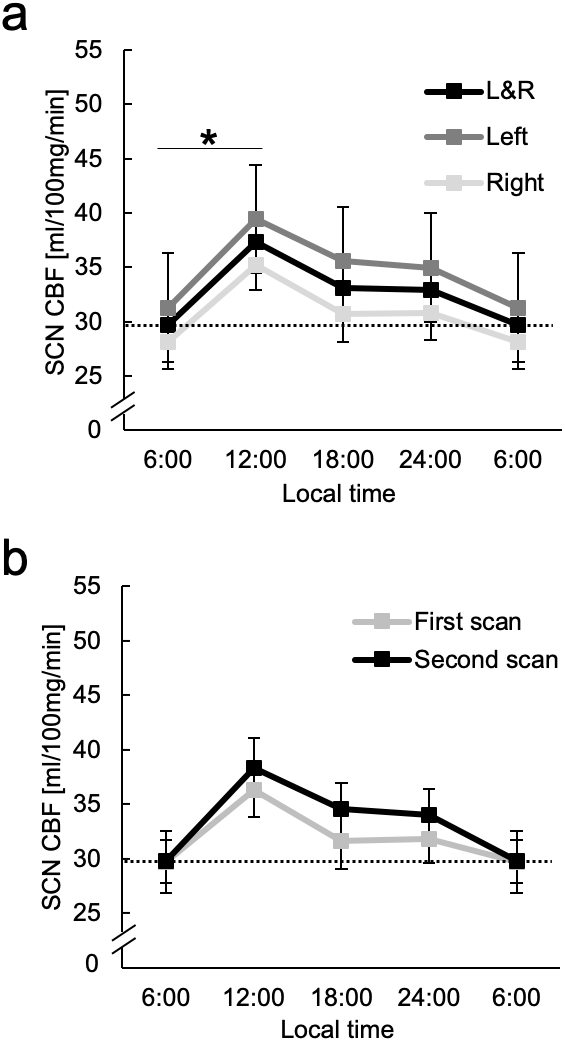
Results of Experiment 1. **a** Time course of the SCN CBF. The CBF signal decreases from the daytime (12:00) to the morning (6:00). The rightmost CBF is the same as the leftmost CBF. The dotted line shows the CBF level at 6:00. **b** Reproducibility of the SCN CBF. The time course from the first scan is similar to that from the second scan. Error bars indicate standard errors of the mean. The asterisk indicates statistical significance (\**P* < 0.05).

A voxel-wise ANOVA around the hypothalamus was performed to examine whether the hypothalamus, other than the SCN, exhibited signal changes. The statistical map clearly showed activity clusters that included the voxels of the SCN and medial preoptic area (MPO) (Supplementary Fig. 3a/b; see also Supplementary Fig. 3c for the time course of MPO activity). The activity pattern was similar to the activity in the adjacent areas of diurnal mammals but not of nocturnal mammals^31–33^.

### SCN activity during the night

In Experiment 2, the SCN activity was measured in detail every 30 min from 24:00 to 6:00. A regression analysis indicates that the SCN activity gradually decreased from midnight to dawn (beta estimate = -1.40, *t*(215) = -2.41, *P* = 0.017) (Fig. 4a).

**Fig. 4.**
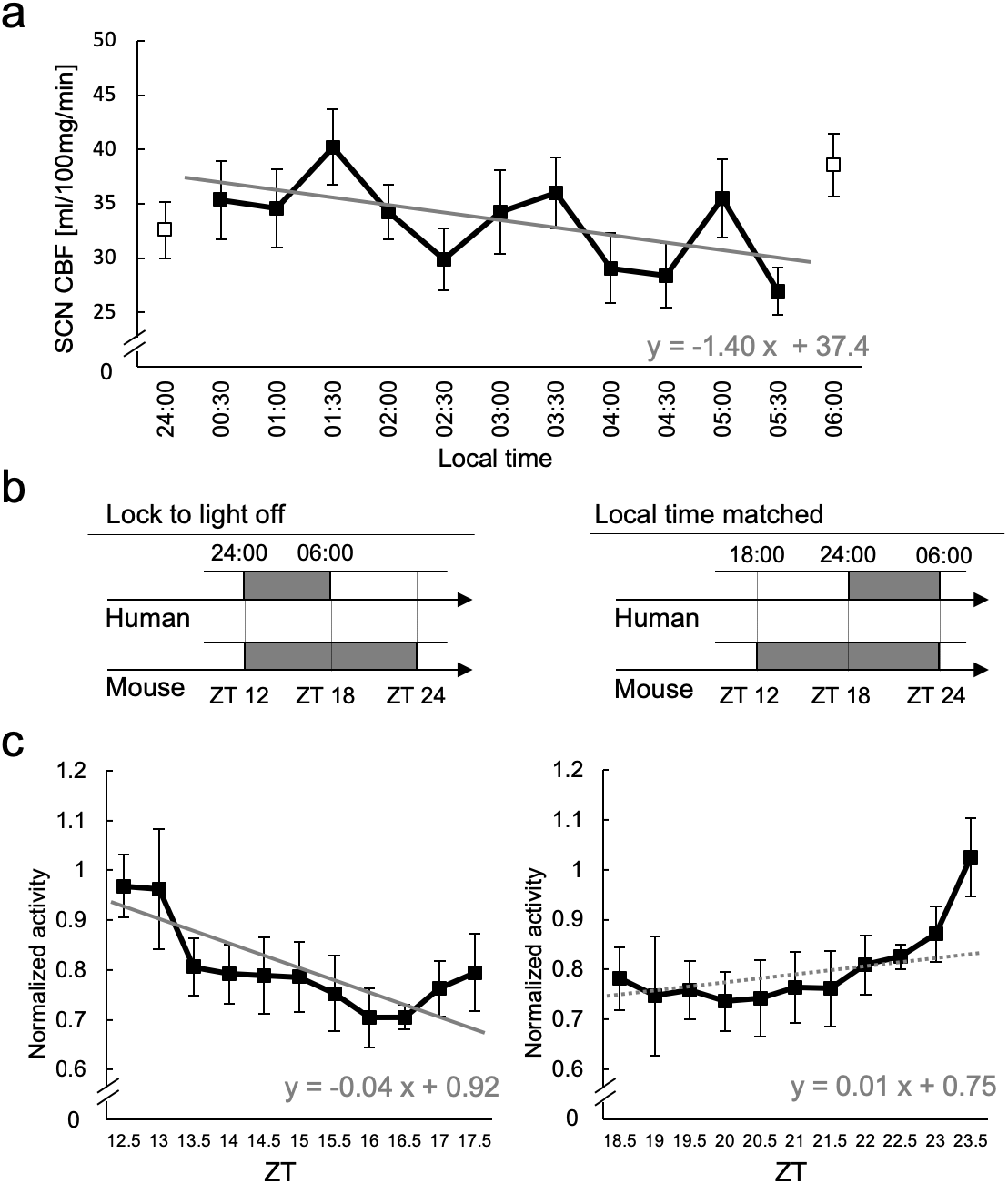
Results of Experiment 2. **a** Time course of the SCN CBF. The white and black dots indicate that the MRI room was light and dark, respectively. The CBF in the SCN decreased gradually during the night. The gray line is a regression line. **b** Two hypotheses of correspondence between the SCN activities of humans and mice. One is the time lock to the switch-off of the room lights. The other is local time matched. **c** Trends of neuronal activity in the rodent SCN. Graphs of the rodent SCN activity were modified from the data between ZT 12 and ZT 18 and between ZT 18 and ZT 24 extracted from Nakamura et al. (2011)^21^. The gray solid and dotted lines are regression lines. Error bars indicate standard errors of the mean.

We then compared the activity of the human SCN with the rodent SCN activity published previously^21^. The previous study is one of very few that collected data, still available to date, from more mature mice that would match in age with the present human study. In the previous study, the multiunit neural activity of the SCN was reported every 30 minutes. The current study divided the rodent SCN data exposed to darkness into two periods: one from zeitgeber time (ZT) 12 to ZT 18 and the other from ZT 18 to ZT 24, where the time of lights on is defined as ZT 0. We examined whether the human SCN activity during the night (24:00 to 6:00) was more compatible with the activity during ZT 12-18 or with the activity during ZT 18-24 in rodents (Fig. 4b). In other words, we examined whether the human SCN activity during the night is more compatible with the rodent SCN activity when the activity is time-locked to lights off or when the activity was matched to local time. The neuronal activity of the rodent SCN decreased during the first half of the dark period (beta estimate = -0.04, *t*(42) = -4.08, *P* = 0.0002) (Fig. 4c left), in accordance with the activity of the human SCN during the night revealed in Experiment 1 at 24:00 and 6:00. In contrast, the activity of the rodent SCN showed an increasing trend during the second half of the dark period (beta estimate = 0.01, *t*(42) = 1.59, *P* = 0.12) (Fig. 4c right). Thus, the activity in the human SCN during the night better matched with the former case in mice.

In order to investigate the relation of the SCN activity to sleep, sleepiness during scanning was reported using a visual analog scale (VAS, from 0 to 10; 10, maximally sleepy or sleeping) after each scan. The temporal effect on the sleepiness VAS score was analyzed using a linear mixed-effects model analysis. The temporal effect was significant for the sleepiness scores (beta estimate = -0.20, *t*(216) = -2.37, *P* = 0.019) (Fig. 5a). Perfusion scans were classified based on the VAS scores of 0 - 10. No significant difference in the SCN activity was observed among the VAS scores (one-way ANOVA, *F*(6, 90) = 0.31, *P* = 0.93) (Fig. 5b), suggesting that the level of sleepiness cannot explain the decreasing trend of the SCN activity during the night. Amylase activity that monitors the stress level measured before and after the MRI session was higher before the experiment (i.e., night) than after the experiment (i.e., morning) (one-tailed t-test, *t*(19) = 1.75, *P* = 0.048), which is consistent with the previous studies^34^, suggesting that the overnight scans were not so stressful to the participants.

**Fig. 5.**
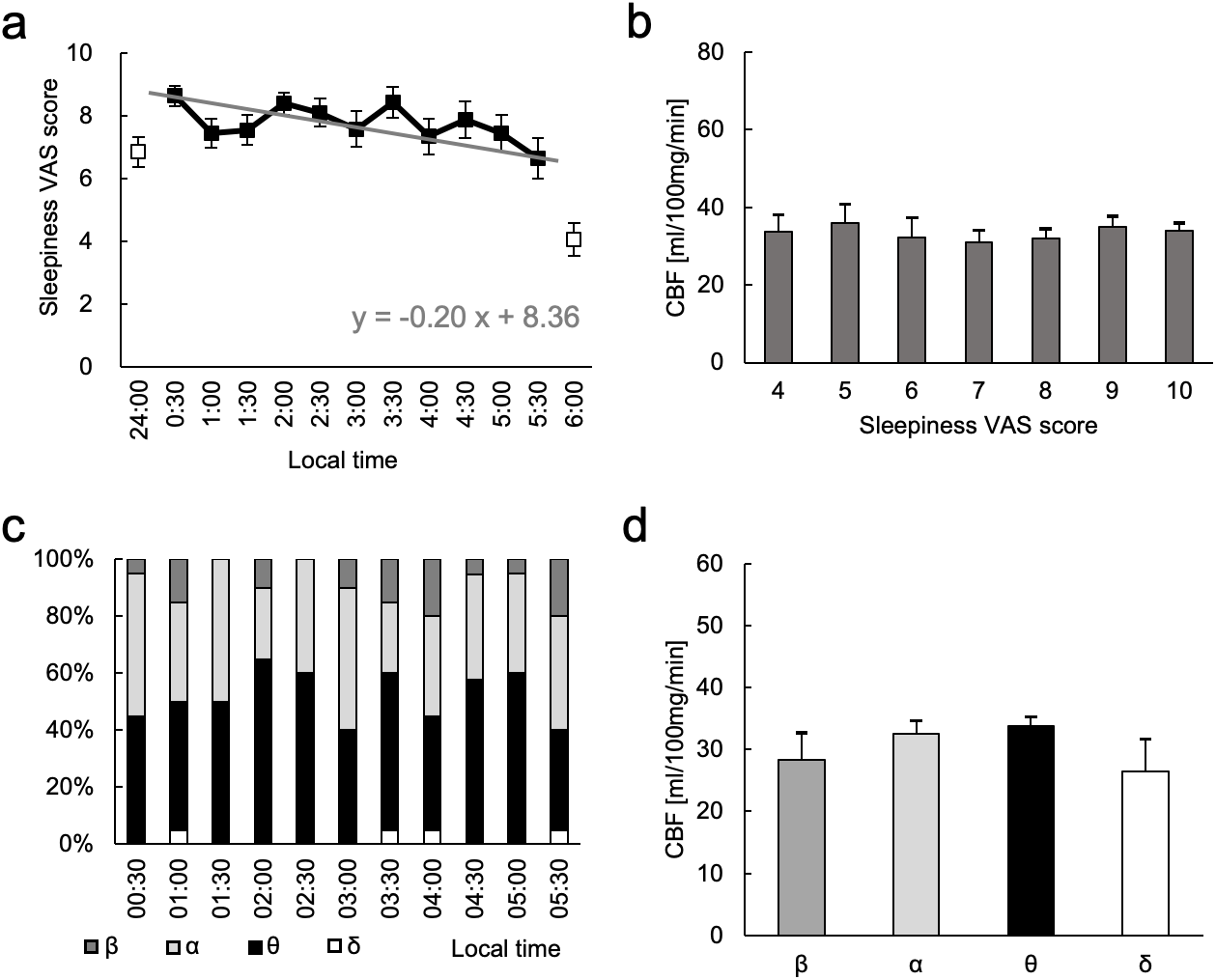
Activity of SCN in Experiment 2. **a** Sleepiness visual-analogue scale (VAS) score. The score decreased during the night. The white dots indicate that the scores were obtained with the lights on in the MRI room, while the black dots indicate that the scores were obtained with the lights off in the MRI room. **b** CBF in the SCN for each sleepiness VAS score. There was no significant difference. **c** Percentages of observed EEG bands in each scan. Alpha and theta bands were mainly observed. **d** SCN CBF of each EEG band. No significant difference between the EEG bands was observed. Error bars indicate standard errors of the mean.

EEG was also recorded during the night scans and classified each perfusion scan into beta/alpha/theta/delta bands. The percentage of EEG frequency band during the first half (before 3:00; beta, 6.0%; alpha, 40.0%; theta, 53.0%; delta 1.0%) was not significantly different from that during the second half (after 3:00; beta 13.1%; alpha, 34.3%; theta, 49.5%; delta 3.0%) (*χ*^2^(3) = 6.49, P = 0.09) (Fig. 5c). Whether SCN activity was different among the EEG bands was also examined. A one-way ANOVA showed that the SCN activity was not significantly different among the EEG bands (*F*(3,46) = 1.17, *P* = 0.33) (Fig. 5d).

With the MPO cluster observed in Experiment 1, additional analyses were conducted. As with the SCN, a regression analysis indicated that the MPO activity showed a gradually decreasing trend from midnight to dawn (beta estimate = -1.35, *t*(215) = -2.64, *P* = 0.0089) (Supplementary Fig. 4a). Sleepiness score was not associated with the MPO activity (one-way ANOVA, *F*(6, 90) = 0.38, *P* = 0.89, Supplementary Fig. 4b), and the activity in the MPO was not significantly different among the EEG bands (one-way ANOVA, *F*(3, 46) = 0.18, *P* = 0.91) (Supplementary Fig. 4c).

## Discussion

This study investigated the diurnal activity of the SCN in humans using perfusion imaging. In the first experiment, the activity of the human SCN was examined every six hours within 24 hours. The activity of the SCN significantly varied, with the maximum activity recorded at noon and the minimum activity at 6:00. The second experiment examined the human SCN activity in more detail during the night (every 30 min from midnight to 6:00), and it was found that the human SCN activity decreased during this period, and the frequency band of the recorded EEG did not significantly affect the SCN activity. Furthermore, the human SCN activity during the night better matched with the activity of the rodent SCN time-locked to lights off. These results suggest that the diurnal variation of the SCN activity in humans was globally consistent with that in non-human mammals and could be influenced locally by lifestyle, such as bright ambient lights late at night.

A boundary mapping technique was employed to allow localization of the SCN in functional images, with a great number of high-resolution (1.25 mm isotropic voxels) BOLD images per participant. The image distortion in the phase-encoding direction around the hypothalamus was corrected using top-up correction, and denoising was conducted using an independent component analysis and machine learning. The coordinates of the SCN voxel in this study were (-2, 2, -16) on the left and (2, 2, -16) on the right, which are very close to those in a recent high-resolution fMRI study^29^: (-5 to - 3, 1 to 4, -17 to -15) on the left and (4 to 6, 1 to 4, -17 to -15) on the right. The slight difference in X coordinates appears to have come from the shape of the third ventricle, which is thinner in coronal slices in our present dataset (Supplementary Fig. 2a/b).

On the other hand, the size of the SCN seems to be less than one voxel of the perfusion imaging (8 mm^3^) of the present study. In a recent cellular-resolution atlas, the size of the SCN was estimated to be (1.7 × 1.1 × 1.1) mm^3^ ∼ 2.1 mm^3^ ^35–37^, and in another recent high-resolution in vivo magnetic resonance imaging atlas, the size was estimated to be 4.9 mm^3^ ^38^. Although the estimated size is still smaller than our 2 mm cubic voxel, the elongated shape of the SCN can be more appropriately covered by a larger ROI, presumably by our 2 mm cubic voxel. Moreover, owing to the 2 mm voxel size larger than the SCN^39^, the location of the SCN voxel was largely appropriate in individual structural images (Supplementary Fig. 2b). It is also to be noted that the areas surrounding the SCN are not classified as nuclei^40^. The estimated distance was 4.5 mm between the SCN and supraoptic nucleus, and 5.5 mm between the SCN and paraventricular nucleus, which is discriminable in the 2 mm voxel/smoothing size. As demonstrated in Supplementary Fig. 2c, it is possible that the adjacent nucleus is included in the 2 mm voxel, but the proportion seems limited.

An increase in neuronal activity leads to metabolic necessity and increases local blood flow in a large part of the brain. However, previous studies have shown that the BOLD signal in the hypothalamus decreased with events such as light exposure, food cue presentation, drinking water, and glucose ingestion^29, 30, 41–43^. Thus, the neurovascular mechanism, which couples the local blood flow and the neural activity, may be inverted in the hypothalamus^44, 45^. The higher blood flow during daytime observed in the present study will not apply to the inverted neurovascular coupling in the hypothalamus; a study of human post-mortem tissues showed the diurnal variations of vasopressin production in the SCN, with higher activity in vasopressin neurons during daytime than during the night^26^. Another example of such ordinary neurovascular coupling in the hypothalamus can be seen in the resting-state functional connectivity between the hypothalamus and the cerebral cortex, which are anatomically connected and yield positive functional connectivity^46–49^. Although precise mechanisms of neurovascular coupling in the hypothalamus remain controversial, it seems reasonable to speculate that neurovascular coupling depends on the nature of the neuronal activity, whether the activity is transiently evoked after an event or spontaneously occurring for a long period.

Neuronal activity in the SCN, which is high during daytime and low during the night, is generated by an endogenous and autonomous circadian rhythm^14, 21, 50–52^. The daily cycle of activity in the SCN is similar in both nocturnal and diurnal mammals^32^. The inverse phase difference of neuronal activity was observed within and outside the SCN in nocturnal mammals^16^. In contrast, the neuronal activity outside the SCN in diurnal mammals showed diurnal variations that paralleled the neuronal activity inside the SCN^31, 53^. Our results showed that the circadian activity in the human SCN was basically similar to that in other diurnal mammals in the SCN and its adjacent areas.

The current study observed a decreasing trend in the SCN activity while the scan room was dark, while increased activity was recorded just after the lights were turned on. Previous animal studies showed that the SCN activity decreased after the lights went out and increased before the lights came on^21, 51, 52, 54, 55^. The decrease in the SCN activity after turning the lights off is consistent for humans and other mammals. On the other hand, the lights in the scan room were turned on at 5:45 in the “one-time” experiment of this study, presumably earlier than the participants’ usual wake-up time. If the experiment is repeated, the rise of the activity in the SCN would shift earlier^54–58^. The human SCN activity during the night may adapt to the period of bright hours in our daily life.

## Materials and Methods

### Experimental designs

In this study, two experiments were conducted, wherein the first investigated the whole cycle of the diurnal activity of the SCN in humans (Experiment 1). Two perfusion images of pseudo-continuous arterial spin labeling (pCASL) in each participant were acquired four times within 24 hours (18:00, 24:00, 6:00, 12:00 on local time) and were used to calculate the CBF at a specific time (Fig. 1a). The lights in the MRI room were turned on in all the scans, and participants were instructed to have meals 4.5 hours before each scan and to rest at a hotel at night (Supplementary Fig. 1). It is a bit unusual to have four meals a day, especially just before sleep at night. Therefore, we instructed the participants to take at least small meals if they do not want to take ordinary meals. To administer scans for multiple (up to four) participants in one 24-h session, the exact scan time was shifted by 30 minutes for each scan, such as 17:00, 17:30, 18:00, and 18:30 for the evening scans.

In the second experiment, we investigated the human SCN activity in more detail during the night (Experiment 2). Participants stayed in the scanner throughout the night, except for brief unavoidable interruptions, and were scanned every 30 min from 24:00 to 6:00 (Fig. 1b). The lights in the MRI room were switched off at 0:15 and on at 5:45 am. In each scan, one single perfusion image was acquired. EEG was recorded during scanning using an MRI-compatible 32-ch international 10/20 EEG system. The level of sleepiness during scanning was reported using VAS (from 0 to 10; 10, maximally sleepy or sleeping) after each scan. To monitor the stress level, salivary amylase activity was measured before and after the MRI session using a salivary amylase monitor (DM-3.1, NIPRO, Osaka, Japan).

### Participants

Twenty-seven right-handed participants without neurological/psychiatric illness or sleep disorders participated in Experiment 1 (13 men and 14 women, age: 22.8 ± 2.7 years [mean ± standard deviation] ranging from 20 to 32 years), while twenty right-handed participants without neurological/psychiatric illness or sleep disorders participated in Experiment 2 (10 men and 10 women, age: 21.5 ± 1.5 years, ranging from 20 to 24 years). None of the participants in these experiments engaged in works that may have affected their circadian rhythms, such as night shifts. Written informed consent was obtained from all participants following the Declaration of Helsinki. Research Ethics Committee, Faculty of Medicine, Juntendo University approved the experimental procedures.

### SCN localization

The location of the SCN was identified using a boundary mapping technique^30, 43, 48, 59–73^. The resting-state images collected by ogawa et al.^30^ were used in this analysis. A total of 3,000 volumes were collected from each of 27 participants with 1.25 mm isotropic voxels using an multiband echo planner imaging (MB-EPI) sequence^74, 75^ (repetition time = 2.3 s, echo time = 20 ms, flip angle = 73°, the in-plane field of view = 180 mm × 180 mm, matrix size = 144 × 144, 108 contiguous slices with no gap, phase encoding direction = posterior-to-anterior, parallel acquisition factor = 2, and multiband factor = 3).

The images were preprocessed using the Human Connectome Project pipeline^76^ with modifications for a higher resolution. The images were motion-corrected, distortion-corrected, and spatially normalized to the standard space of Montreal Neurological Institute (MNI) coordinates. The time series of the images were projected from the voxel space onto the standard surface (32,492 vertices in each hemisphere)^76^, high-pass filtered (cut-off = 2000 s), and denoised using the ICAFIX method^77^. Surface registration was refined using the MSM-All^78^. Global signals were regressed out using the mean time series across the surface vertices.

We calculated correlations between each voxel in the hypothalamus, excluding the voxels in the mammillary body, and the vertices in the cerebral surface of each participant. The correlation coefficient in each vertex was transformed to Fisher’s z value (i.e., cerebral correlation map), and the spatial similarity of the cerebral correlation maps for hypothalamic voxels was then calculated (i.e., hypothalamic similarity map). Spatial gradients of the hypothalamic similarity maps were computed for each hypothalamic voxel (i.e., hypothalamic gradient map). The hypothalamic gradient maps were averaged across participants to generate the group hypothalamic gradient maps. After a minimal spatial smoothing (full width at half-maximum [FWHM] = 1.25 mm) of the group hypothalamic gradient maps, excluding the third ventricle of each side, a three-dimensional watershed algorithm^79^ was applied to the group hypothalamic gradient maps (i.e., binary watershed maps). The binary watershed maps were averaged across the hypothalamic voxels to generate the probabilistic boundary map (Fig. 1c, 1d), where the probability value represents how likely the voxel is a boundary of the functional area (parcel), or in other words, how unlikely the voxel is a center of the parcel.

The bilateral parcels of the SCN were identified above the optic chiasm, and the center voxel of the SCN parcel was defined as the voxel with the least probability value in the SCN parcel in a 1.25 mm isotropic BOLD image. The SCN ROI in a 2 mm isotropic perfusion image was defined as the voxel corresponding to the center voxel in the SCN parcel. Therefore, the volume of the SCN ROI was 8 mm^3^, consisting of one 2 mm isotropic voxel.

### MRI procedures

All MRI data were acquired using a 3-T MRI scanner at Juntendo University Hospital (Siemens Prisma, Erlangen, Germany) with a 32-ch head coil. T1- weighted structural images were obtained using 3D magnetization-prepared rapid gradient-echo (resolution = 0.8 × 0.8 × 0.8 mm^3^). Whole brain perfusion images were acquired using pCASL imaging with MB-EPI^80^ (number of measurements = 90, repetition time = 4.0 s, echo time = 25.2 ms, partial Fourier = 6/8, flip angle = 90°, labeling duration = 1.5 s, post labeling delay = 1.64 s, slice thickness = 1.82 mm, distance factor = 10%, number of slices = 72, slice acquisition order = ascending, in-plane resolution = 106 × 106 mm^2^, multiband acceleration factor = 6). Two images with one anterior-to-posterior and one posterior-to-anterior encoding direction were acquired using the spin-echo field map sequence before each pCASL imaging. These images were used to perform the top-up distortion correction for pCASL images^81^.

The perfusion images were corrected for motion and distortion. The CBF maps in the standard MNI space were calculated using a command line interface of oxford_asl^82^. Spatially minimal smoothing was applied to the CBF images (full-wise half maximum of Gaussian kernel = 2.0 mm), and the CBF values of the SCN and MPO ROIs were extracted and sent for statistical analyses.

### EEG procedures

In Experiment 2, EEG was recorded during the scan using an MRI-compatible 32-ch international 10/20 EEG system (Geodesic Sensor Net, Electrical Geodesics, Inc., Eugene, Oregon, USA). Before the experiment, the impedances of all electrodes were adjusted to < 50 kΩ. The impedances were checked before each scan. After the experiment, EEG data were band-pass filtered (0.3-30 Hz) and referenced to the electrode Cz. The EEG data around the middle of each scan was used for the classification of the frequency band: beta (more than 12 Hz), alpha (8-12 Hz), theta (4-8 Hz), or delta (1-4 Hz) band. One of the authors classified the frequency band of EEG data of each scan, and then the other author verified the classification. The data of electrodes whose impedance was over 50 kΩ in any scan were not used for the EEG band classification.

## Acknowledgments

This work was supported by JSPS KAKENHI Grants 22K07334 to A.O. and 21K07255 to T.O., a grant from Takeda Science Foundation to S.K, and a Grant-in-Aid for Special Research in Subsidies for ordinary expenses of private schools from The Promotion and Mutual Aid Corporation for Private Schools of Japan.

## Data and code availability

The data and code that support the findings of this study are available at Dryad, except for raw image data. The raw image data cannot be deposited in a public repository because sharing raw image data was not included in the informed consent. Any additional information required to reanalyze the data reported in this paper is available from the corresponding author upon reasonable request.

---

**Supplementary Fig. 1.**
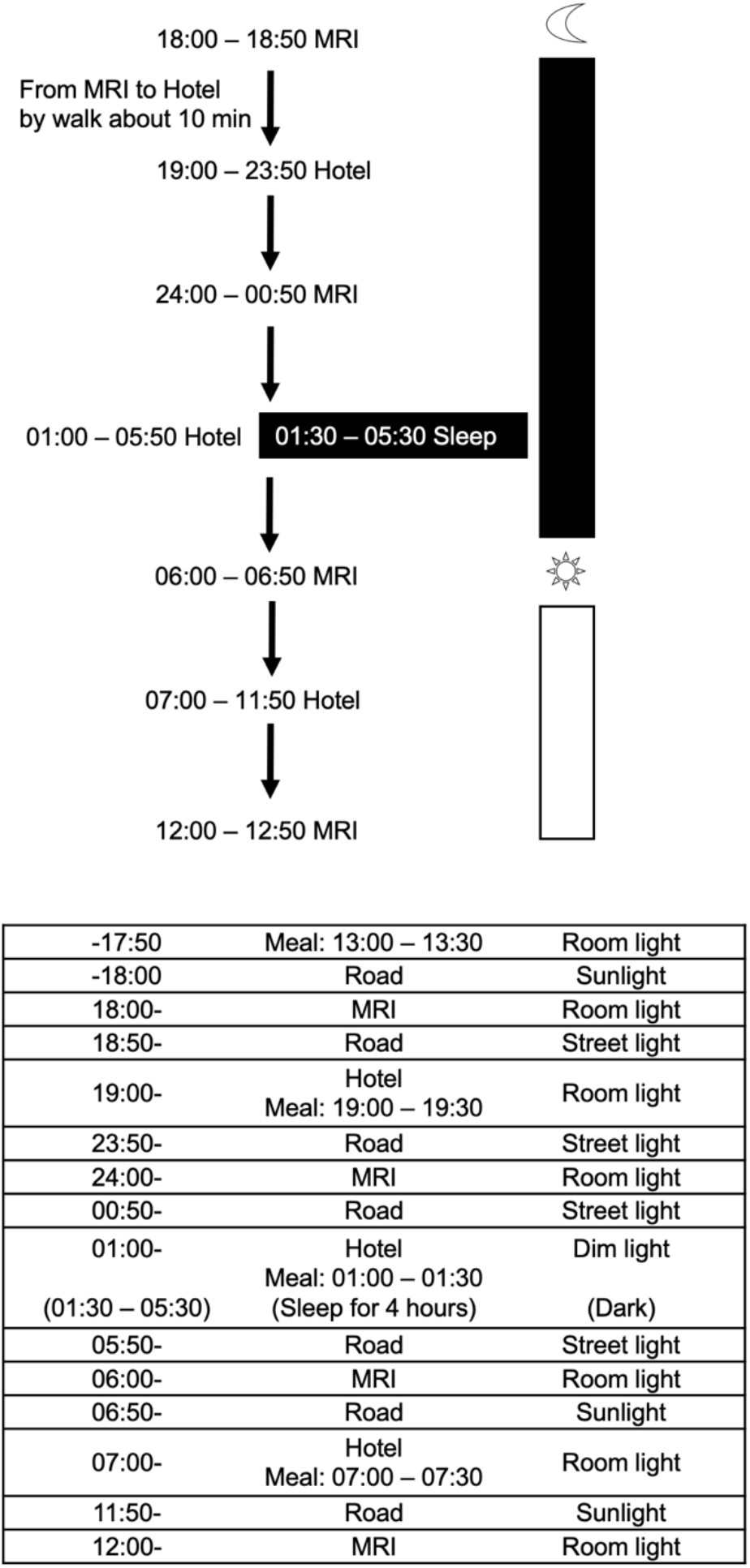
Example of schedule in Experiment 1. Participants took a rest at a hotel between MRI scans.

**Supplementary Fig. 2.**
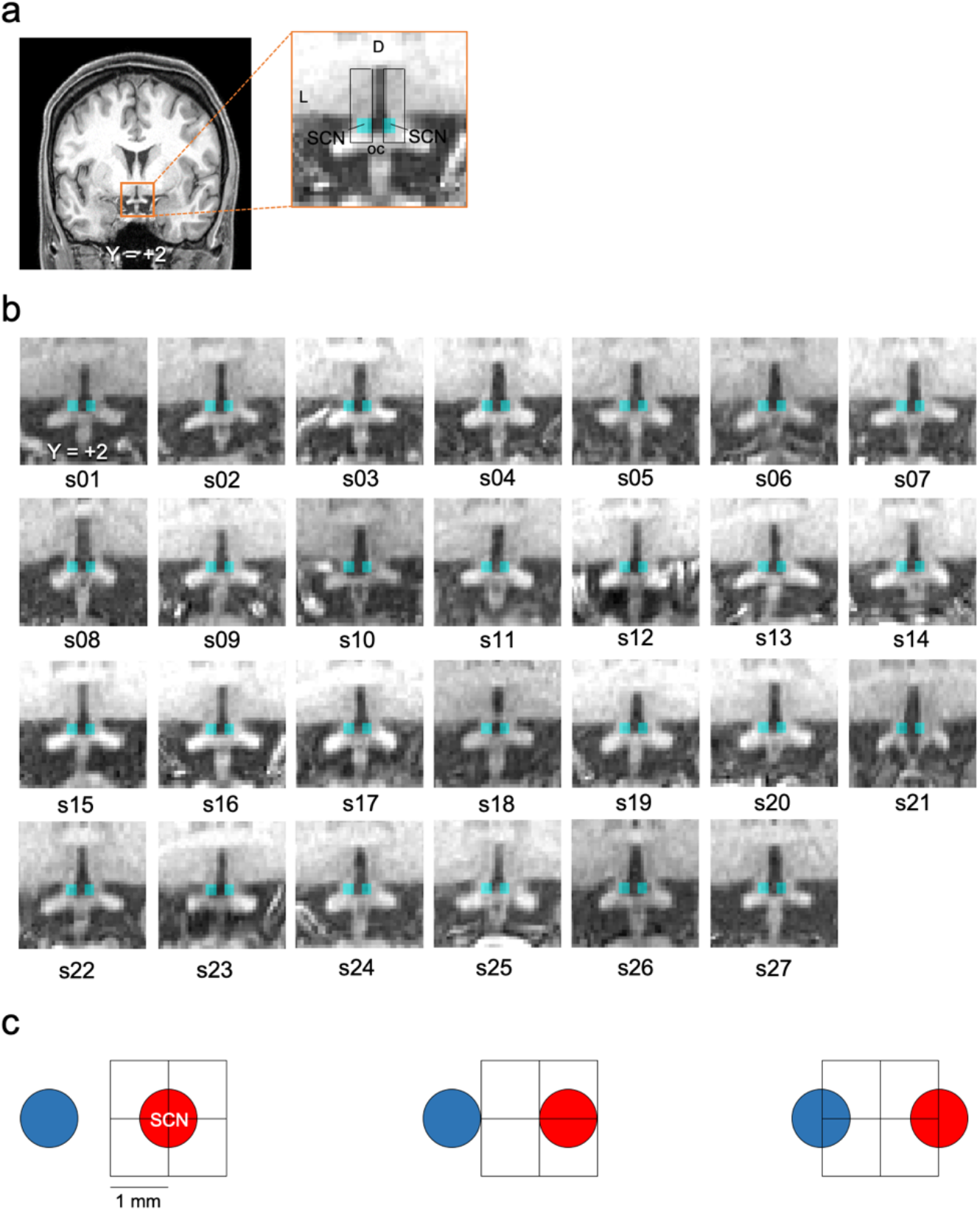
SCN ROIs. **a** SCN ROIs on the coronal section of a high-resolution T1-weighted image of a representative participant. The cyan voxels indicate the SCN ROIs. **b** SCN ROIs on the coronal sections of high-resolution T1-weighted images of all participants. **c** Spatial coverage of the SCN using a 2-mm isotropic voxel in relation to another nucleus. In the left panel, when the SCN (approximately 1 mm diameter) is located in the center of the 2 mm voxel, it is unlikely that the adjacent nucleus (approximately 1 mm diameter) located approximately 1 mm apart is included in that voxel. In the middle panel, when the SCN is located 0.5 mm off the center of the voxel, only a small portion of the adjacent nucleus would be included in that voxel. In the right panel, when the SCN is not fully included in the voxel, the adjacent nucleus may be included to an inverse degree. Red and blue disks indicate the SCN and another nucleus of the hypothalamus, such as the supraoptic nucleus. The scale bar indicates 1 mm.

**Supplementary Fig. 3.**
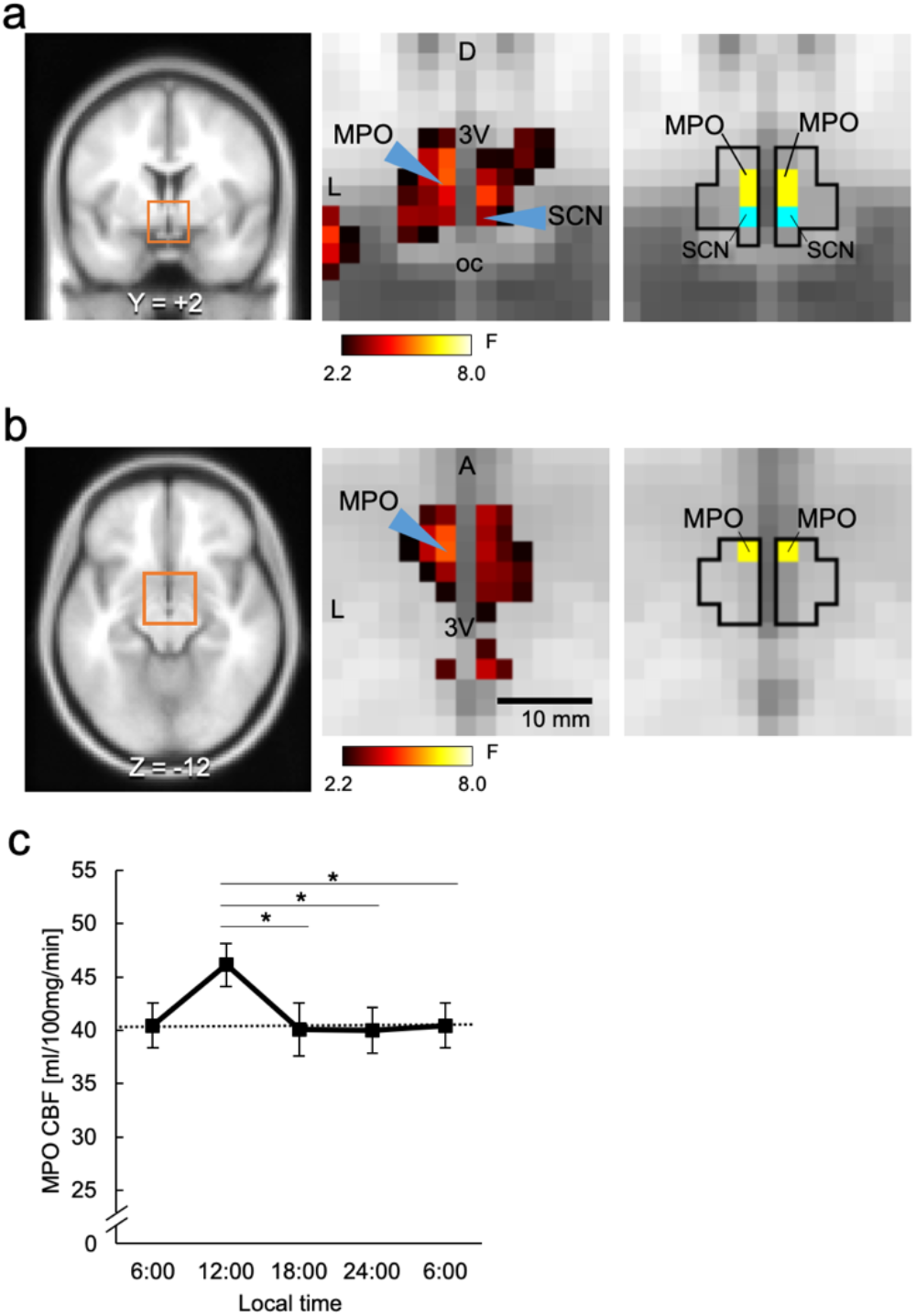
Statistical map around the hypothalamus, ROI of MPO, and time course of MPO CBF. **a** F-map and the MPO position on the coronal section. The black solid lines in the right panel indicate the border of the hypothalamus. The cyan and yellow voxels indicate ROIs of the SCN and MPO, respectively. **b** F-map and the MPO position on the axial section. The black solid lines in the right panel indicate the border of the hypothalamus. The coordinates of MPO ROI were x = -2, y = 2, z = - 14 to -12 for the left and x = 2, y = 2, z = -14 to -12 for the right. **c** Time course of MPO CBF. There was a significant temporal effect (one-way repeated-measures ANOVA, *F*(3,78) = 4.35, *P* = 0.0069). The dot-line indicates the CBF at 6:00. CBF at 12:00 was significantly higher than those at 18:00, 24:00, and 6:00. Error bars indicate standard errors of the mean. The asterisks indicate statistical significance (\**P* < 0.05).

**Supplementary Fig. 4.**
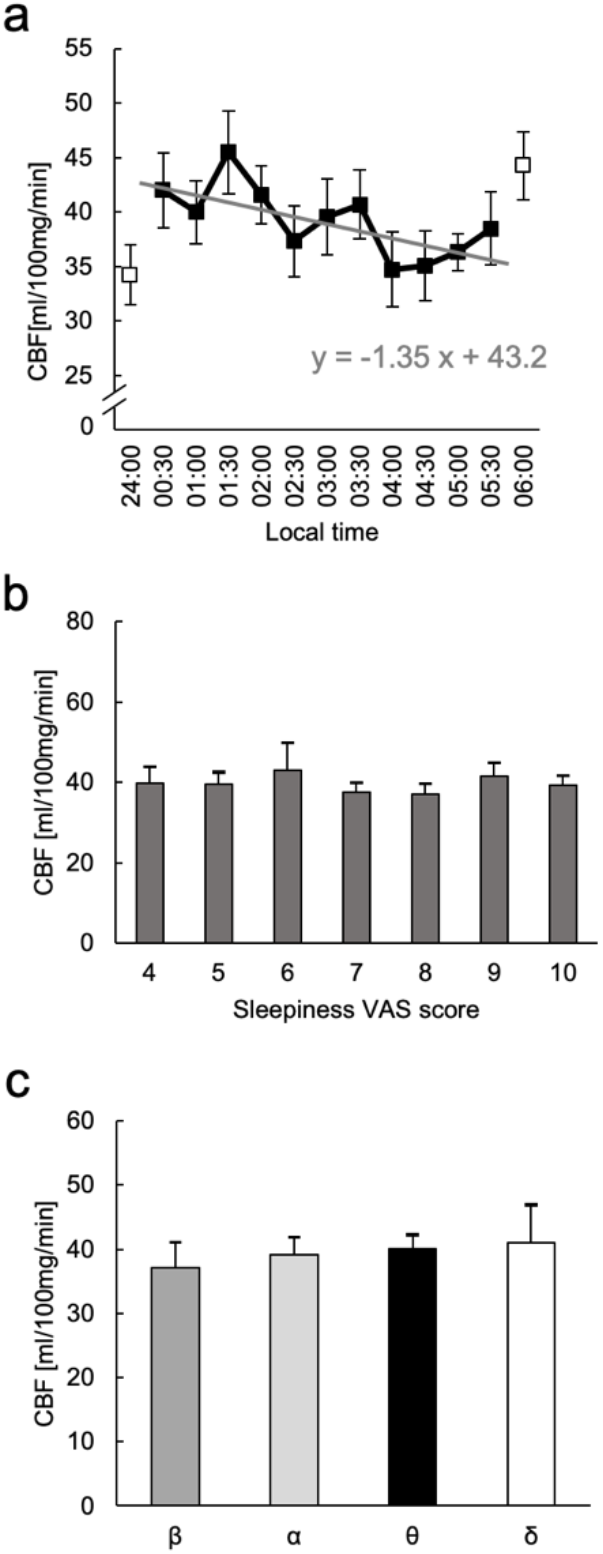
Supplementary results of Experiment 2. **a** Time course of CBF in MPO. The white and black dots indicate that the light of MRI room was on and off, respectively. **b** CBF in MPO for each sleepiness VAS score. As with SCN, no significant difference was observed. **c** CBF in MPO in each EEG band. There is no significant difference between the EEG bands. Error bars indicate standard errors of the mean.

## References

1. Hastings, M. H., Maywood, E. S. & Brancaccio, M. Generation of circadian rhythms in the suprachiasmatic nucleus. Nat. Rev. Neurosci. 19, 453–469 (2018).

2. Cai, X. et al. Imaging the effect of the circadian light-dark cycle on the glymphatic system in awake rats. Proc. Natl. Acad. Sci. 117, 668–676 (2020).

3. Fishbein, A. B., Knutson, K. L. & Zee, P. C. Circadian disruption and human health. J. Clin. Investig. 131, (2021).

4. Moore, R. Y. & Eichler, V. B. Loss of a circadian adrenal corticosterone rhythm following suprachiasmatic lesions in the rat. Brain Res. 42, 201–206 (1972).

5. Stephan, F. K. & Zucker, I. Circadian Rhythms in Drinking Behavior and Locomotor Activity of Rats Are Eliminated by Hypothalamic Lesions. Proc. Natl. Acad. Sci. 69, 1583–1586 (1972).

6. Sumova, A., Travnicnkova&, Z., Peterst, R., Schwartzt, W. J. & Illnerova, H. The rat suprachiasmatic nucleus is a clock for all seasons. Proc. Natl. Acad. Sci. 92, 7754–7758 (1995).

7. Nagano, M. et al. An Abrupt Shift in the Day/Night Cycle Causes Desynchrony in the Mammalian Circadian Center. J. Neurosci. 23, 6141–6151 (2003).

8. Enoki, R. et al. Topological specificity and hierarchical network of the circadian calcium rhythm in the suprachiasmatic nucleus. Proc. Natl. Acad. Sci. 109, 21498–21503 (2012).

9. Yamaguchi, S. et al. Synchronization of Cellular Clocks in the Suprachiasmatic Nucleus. Science (1979) 302, 1408–1412 (2003).

10. Welsh, D. K., Takahashi, J. S. & Kay, S. A. Suprachiasmatic Nucleus: Cell Autonomy and Network Properties. Annu. Rev. Physiol. 72, 551–577 (2010).

11. Welsh, D. K., Logothetis, D. E., Meister, M. & Reppert, S. M. Individual Neurons Dissociated from Rat Suprachiasmatic Nucleus Express Independently Phased Circadian Firing Rhythms. Neuron 14, 697–706 (1995).

12. Saper, C. B., Lu, J., Chou, T. C. & Gooley, J. The hypothalamic integrator for circadian rhythms. Trends Neurosci. 28, 152–157 (2005).

13. Saper, C. B., Scammell, T. E. & Lu, J. Hypothalamic regulation of sleep and circadian rhythms. Nature 437, 1257–1263 (2005).

14. Inouye, S. T. & Kawamura, H. Persistence of circadian rhythmicity in a mammalian hypothalamic ‘island’ containing the suprachiasmatic nucleus. Proc. Natl. Acad. Sci. 76, 5962–5966 (1979).

15. Inouye, S. T. & Kawamura, H. Characteristics of a circadian pacemaker in the suprachiasmatic nucleus. J. Comp. Physiol. 146, 153–160 (1982).

16. Musiek, E. S. & Holtzman, D. M. Mechanisms linking circadian clocks, sleep, and neurodegeneration. Science 354, 1004–1008 (2016).

17. Hattar, S., Liao, H.-W., Takao, M., Berson, D. M. & Yau, K.-W. Melanopsin- Containing Retinal Ganglion Cells: Architecture, Projections, and Intrinsic Photosensitivity. Science 295, 1065–1070 (2002).

18. Fernandez, D. C. et al. Light Affects Mood and Learning through Distinct Retina-Brain Pathways. Cell 175, 71–84.e18 (2018).

19. Berson, D. M., Dunn, F. A. & Takao, M. Phototransduction by Retinal Ganglion Cells That Set the Circadian Clock. Science 295, 1070–1073 (2002).

20. Touitou, Y. & Point, S. Effects and mechanisms of action of light-emitting diodes on the human retina and internal clock. Environ. Res. 190, 109942 (2020).

21. Nakamura, T. J. et al. Age-related decline in circadian output. J. Neurosci. 31, 10201–10205 (2011).

22. Vandewalle, G. et al. Spectral quality of light modulates emotional brain responses in humans. Proc. Natl. Acad. Sci. 107, 19549–19554 (2010).

23. Legates, T. A. et al. Aberrant light directly impairs mood and learning through melanopsin-expressing neurons. Nature 491, 594–598 (2012).

24. Fernandez, F. et al. Dysrhythmia in the suprachiasmatic nucleus inhibits memory processing. Science 346, 854–857 (2014).

25. Ruby, N. F. et al. Hippocampal-dependent learning requires a functional circadian system. Proc. Natl. Acad. Sci. 105, 15593–15598 (2008).

26. Hofman, M. A. & Swaab, D. F. Alterations in circadian rhythmicity of the vasopressin-producing neurons of the human suprachiasmatic nucleus (SCN) with aging. Brain Res. 651, 134–142 (1994).

27. Hofman, M. A. & Swaab, D. F. Diurnal and seasonal rhythms of neuronal activity in the suprachiasmatic nucleus of humans. J. Biol. Rhythms 8, 283–295 (1993).

28. Hofman, M. A. The human circadian clock and aging. Chronobiol. Int. 17, 245– 259 (2000).

29. Schoonderwoerd, R. A., et al. The photobiology of the human circadian clock. Proc. Natl. Acad. Sci. 119, e21e2118803119 (2022).

30. Ogawa, A. et al. Hypothalamic interaction with reward-related regions during subjective evaluation of foods. Neuroimage 264, 119744 (2022).

31. Sato, T. & Kawamura, H. Circadian rhythms in multiple unit activity inside and outside the suprachiasmatic nucleus in the diurnal chipmunk (Eutamias sibiricus). Neurosci. Res. 1, 45–52 (1984).

32. Bano-Otalora, B. et al. Daily electrical activity in the master circadian clock of a diurnal mammal. Elife 10, e68179 (2021).

33. Bano-Otalora, B., et al. Bright daytime light enhances circadian amplitude in a diurnal mammal. Proc. Natl. Acad. Sci. 118, e2100094118 (2021).

34. Nater, U. M., Rohleder, N., Schlotz, W., Ehlert, U. & Kirschbaum, C. Determinants of the diurnal course of salivary alpha-amylase. Psychoneuroendocrinology 32, 392–401 (2007).

35. Vimal, R. L. P. et al. Activation of suprachiasmatic nuclei and primary visual cortex depends upon time of day. Eur. J. Neurosci. 29, 399–410 (2009).

36. Sharifpour, R. et al. Pitfalls in recording BOLD signal responses to light in small hypothalamic nuclei using Ultra-High-Field 7 Tesla MRI. Proc. Natl. Acad. Sci. 119, (2022).

37. Ding, S. et al. Comprehensive cellular-resolution atlas of the adult human brain. J. Comp. Neurol. 524, 3127–3481 (2016).

38. Neudorfer, C. et al. A high-resolution in vivo magnetic resonance imaging atlas of the human hypothalamic region. Sci. Data 7, 305 (2020).

39. Meijer, J. H., et al. Reply to Sharifpour, et al.: Light response measurement of the human SCN by 7T fMRI. Proc. Natl. Acad. Sci. 119, (2022).

40. Mai, J., Majtanik, M. & Paxinos, G. Atlas of the Human Brain 4th Edition. (Academic Press, 2015).

41. Smeets, P. A. M., de Graaf, C., Stafleu, A., van Osch, M. J. P. & van der Grond, J. Functional MRI of human hypothalamic responses following glucose ingestion. Neuroimage 24, 363–368 (2005).

42. Vidarsdottir, S. et al. Glucose ingestion fails to inhibit hypothalamic neuronal activity in patients with type 2 diabetes. Diabetes 56, 2547–2550 (2007).

43. Osada, T. et al. Functional subdivisions of the hypothalamus using areal parcellation and their signal changes related to glucose metabolism. Neuroimage 162, 1–12 (2017).

44. Roy, R. K. et al. Inverse neurovascular coupling contributes to positive feedback excitation of vasopressin neurons during a systemic homeostatic challenge. Cell Rep. 37, 109925 (2021).

45. Drew, P. J. Neurovascular coupling: motive unknown. Trends Neurosci. 45, 809–819 (2022).

46. Kullmann, S. et al. Resting-state functional connectivity of the human hypothalamus. Hum. Brain Mapp. 35, 6088–6096 (2014).

47. Hirose, S. et al. Lateral–Medial Dissociation in Orbitofrontal Cortex– Hypothalamus Connectivity. Front. Hum. Neurosci. 10, 244 (2016).

48. Ogawa, A. et al. Connectivity-based localization of human hypothalamic nuclei in functional images of standard voxel size. Neuroimage 221, 117205 (2020).

49. Zhang, S., Wang, W., Zhornitsky, S. & Li, C. S. R. Resting state functional connectivity of the lateral and medial hypothalamus in cocaine dependence: An exploratory study. Front. Psychiatry 9, 344 (2018).

50. Kuhlman, S. J. & McMahon, D. G. Encoding the ins and outs of circadian pacemaking. J. Biol. Rhythms 21, 470–481 (2006).

51. Nakamura, W. et al. In Vivo Monitoring of circadian timing in freely moving mice. Current Biology 18, 381–385 (2008).

52. Miyamoto, H., Nakamaru-Ogiso, E., Hamada, K. & Hensch, T. K. Serotonergic integration of circadian clock and ultradian sleep-wake cycles. J. Neurosci. 32, 14794–14803 (2012).

53. Li, J. Z. et al. Circadian patterns of gene expression in the human brain and disruption in major depressive disorder. Proc. Natl. Acad. Sci. 110, 9950–9955 (2013).

54. Schaap, J. et al. Heterogeneity of rhythmic suprachiasmatic nucleus neurons: Implications for circadian waveform and photoperiodic encoding. Proc. Natl. Acad. Sci. 100, 15994–15999 (2003).

55. VanderLeest, H. T. et al. Seasonal encoding by the circadian pacemaker of the SCN. Current Biology 17, 468–473 (2007).

56. Brown, T. M. & Piggins, H. D. Spatiotemporal heterogeneity in the electrical activity of suprachiasmatic nuclei neurons and their response to photoperiod. J. Biol. Rhythms 24, 44–54 (2009).

57. Houben, T., Deboer, T., van Oosterhout, F. & Meijer, J. H. Correlation with behavioral activity and rest implies circadian regulation by SCN neuronal activity levels. J. Biol. Rhythms 24, 477–487 (2009).

58. VanderLeest, H. T., Rohling, J. H. T., Michel, S. & Meijer, J. H. Phase shifting capacity of the circadian pacemaker determined by the SCN neuronal network organization. PLoS ONE 4, (2009).

59. Suda, A. et al. Functional organization for response inhibition in the right inferior frontal cortex of individual human brains. Cereb. Cortex 30, 6325–6335 (2020).

60. Nakajima, K. et al. A causal role of anterior prefrontal-putamen circuit for response inhibition revealed by transcranial ultrasound stimulation in humans. Cell Rep. 40, 111197 (2022).

61. Osada, T. et al. Parallel cognitive processing streams in the human prefrontal cortex: Parsing the areal-level brain network for response inhibition. Cell Rep. 36, 109732 (2021).

62. Margulies, D. S. et al. Mapping the functional connectivity of anterior cingulate cortex. Neuroimage 37, 579–588 (2007).

63. Cohen, A. L. et al. Defining functional areas in individual human brains using resting functional connectivity MRI. Neuroimage 41, 45–57 (2008).

64. Buckner, R. L., Krienen, F. M., Castellanos, A., Diaz, J. C. & Yeo, B. T. T. The organization of the human cerebellum estimated by intrinsic functional connectivity. J. Neurophysiol. 106, 2322–45 (2011).

65. Eickhoff, S. B., Thirion, B., Varoquaux, G. & Bzdok, D. Connectivity-based parcellation: Critique and implications. Hum. Brain Mapp. 36, 4771–4792 (2015).

66. Laumann, T. O. et al. Functional system and areal organization of a highly sampled individual human brain. Neuron 87, 1–14 (2015).

67. Poldrack, R. A. et al. Long-term neural and physiological phenotyping of a single human. Nat. Commun. 6, 8885 (2015).

68. Glasser, M. F. et al. A multi-modal parcellation of human cerebral cortex. Nature 536, 171–178 (2016).

69. Gordon, E. M. et al. Generation and evaluation of a cortical area parcellation from resting-state correlations. Cereb. Cortex 26, 288–303 (2016).

70. Gratton, C. et al. Functional brain networks are dominated by stable group and individual factors, not cognitive or daily variation. Neuron 98, 439–452.e5 (2018).

71. Ogawa, A. et al. Striatal subdivisions that coherently interact with multiple cerebrocortical networks. Hum. Brain Mapp. 39, 4349–4359 (2018).

72. Fujimoto, U. et al. Network centrality analysis characterizes brain activity during response inhibition in right ventral inferior frontal cortex. Juntendo Medical Journal 68, 208–211 (2022).

73. Fujimoto, U. et al. Network centrality reveals dissociable brain activity during response inhibition in human right ventral part of inferior frontal cortex. Neuroscience 433, 163–173 (2020).

74. Xu, J. et al. Evaluation of slice accelerations using multiband echo planar imaging at 3T. Neuroimage 83, 991–1001 (2013).

75. Feinberg, D. A. et al. Multiplexed echo planar imaging for sub-second whole brain fmri and fast diffusion imaging. PLoS ONE 5, e15710 (2010).

76. Glasser, M. F. et al. The minimal preprocessing pipelines for the Human Connectome Project. Neuroimage 80, 105–124 (2013).

77. Salimi-Khorshidi, G. et al. Automatic denoising of functional MRI data: Combining independent component analysis and hierarchical fusion of classifiers. Neuroimage 90, 449–468 (2014).

78. Robinson, E. C. et al. Multimodal surface matching with higher-order smoothness constraints. Neuroimage 167, 453–465 (2018).

79. Vincent, L. & Soille, P. Watersheds in digital spaces: An efficient algorithm based on immersion simulations. IEEE Trans. Pattern Anal. Mach. Intell. 13, 583–598 (1991).

80. Li, X. et al. Theoretical and experimental evaluation of multi-band EPI for high- resolution whole brain pCASL imaging. Neuroimage 106, 170–181 (2015).

81. Andersson, J. L. R., Skare, S. & Ashburner, J. How to correct susceptibility distortions in spin-echo echo-planar images: Application to diffusion tensor imaging. Neuroimage 20, 870–888 (2003).

82. Chappell, M. A., Groves, A. R., Whitcher, B. & Woolrich, M. W. Variational Bayesian Inference for a Nonlinear Forward Model. IEEE Trans. Signal Process. 57, 223–236 (2009).

